# Transcriptional Changes of DNA Replication and Repair Factors Over Uveal Melanoma Subtypes

**DOI:** 10.1101/214932

**Authors:** Melanie Kucherlapati

## Abstract

**Background:** Uncontrolled replication is a process common to all cancers facilitated by the summation of changes accumulated as tumors progress. The aim of this study was to examine small groups of genes with known biology in replication and repair at the transcriptional and genomic levels, correlating alterations with survival in Uveal Melanoma tumor progression. Selected components of Pre-Replication, Pre-Initiation, and Replisome Complexes, DNA Damage Response and Mismatch Repair have been observed.

**Methods:** We have generated two groups for each gene examined above and below the average alteration level, and compared relative expression and survival across TCGA UVM subtypes based on somatic copy number alteration supported by DNA methylation and mRNA/miRNA/lncRNA expression. Significance between subtypes monosomic or disomic for chromosome 3 was determined by Fisher’s exact test. Kaplan Meier survival distribution based on disease specific survival was compared by log-rank test.

**Results:** Specific genes with significant alteration include MCM2 MCM4 and MCM5 of the Minichromosome Maintenance helicase complex, CDC45, MCM10, CIZ1, PCNA, FEN1, LIG1, POLD1, POLE, HUS1, CHECK1, ATRIP, MLH3, and MSH6. We found evidence of Exon 4 skipping in CIZ1 previously identified as a cancer variant and reportedly used as an early serum biomarker in lung cancer, accompanied by evidence of instability of a mononucleotide repeat in Intron 3. Mismatch Repair protein MLH3 was found to have splicing variations with deletions to both Exon 5 and Exon 7 simultaneously. PCNA, FEN1, and LIG1 had increased relative expression levels not due to their mutation or to copy number variation.

**Conclusion:** We have observed differences in relative and differential expression that support the concept that selected replication and repair genes and their products are causally involved in the origin and progression of uveal melanoma, suggesting specific avenues for early biomarker identification and also therapeutic approach.

## Background

Comparatively uncontrolled replication carried out by highly evolutionarily conserved multiprotein complexes, is a process shared by all cancers. Although many other processes including immortality, epithelial to mesenchyme transition, telomere metabolism, metastasis etc. contribute to tumorigenesis, the summation of genomic alterations in a tumor must facilitate replication. Duplication in the transformed cell is achieved at the expense of decreased fidelity, making replication a focal point where heterogeneity is created that upon clonal selection leads to tumor expansion and survival. The status of several individual critical replication genes has been examined in The Cancer Genome Atlas (TCGA) efforts and auxiliary studies, with computational methods placing alterations into known pathways to identify potential targets for precision medicine. However, the behavior of replication factors as a group did not receive analogous systematic investigation, likely because they are thought of as being part of a process rather than a pathway.

In this study we have examined the status of small groups of genes with known biology in replication and repair at the genomic and transcriptional levels in one tumor type, Uveal Melanoma (UVM). TCGA has recently conducted an integrative analysis of this cancer, and the data providing its molecular underpinnings are now available to the public as part of the “Rare Tumor Project” [1]. While having a low incidence of 5.1 per million [2], UVM is the most common intraocular malignancy. It is highly lethal, with 50% of patients developing metastatic disease followed by 6–12 month survival from metastatic diagnosis [3]. There are no effective treatments for metastatic UVM, early diagnosis could lower lethality.

TCGA has examined this tumor type and generated datasets through whole exome sequencing (WES), whole genome sequencing (WGS), mRNA miRNA and lncRNA expression, DNA methylation, identification of immune infiltration, and detailed pathology with clinical outcome. Based on the combination of somatic copy number alterations (SCNA), DNA methylation patterns, mRNA miRNA and lncRNA expression, tumors were classified into four unsupervised clusters designated 1 through 4. Pathology and clinical data shows increasing malignancy across the clusters, with subtypes 3 and 4 having poorer outcomes that included shorter time to metastasis.

Selected components from the Pre-replication, Pre-initiation, Replisome, DNA Damage Repair (DDR), and Mismatch Repair (MMR) complexes have been investigated in this study for genomic and relative transcriptional changes across the UVM clusters. We have found alterations indicative of replication stress correlating with aneuploidy, increased malignancy, and decreased survival. Replication stress is thought to be an early and strong driving force in tumorigenesis. It is defined as impediment to the replication fork that causes slowing or stalling of the replication machinery and is brought on by a variety of factors broadly classified as “exogenous” and “endogenous” [4]. Examples of exogenous causes are radiation, therapeutic treatment, and diet, where endogenous causes are DNA structures, protein-DNA complexes, nucleotide pool availability, reactive oxidation species, transcription and replication complex collision, and mutation and expression alteration in tumor suppressor and oncogenes. Replication stress is common in cancer and has been previously suggested as therapeutically targetable [5, 6]. Common components integral to replication involved in error prone bypass of damage are also likely effective points of intervention [7, 8].

In a normal cell replication stress will activate DDR which prevents DNA damage from becoming fixed during replication and passed on in mitosis [9]. It does this in part by coordinating cell cycle control and providing a pause for repair, and also in some cases by triggering apoptosis. Dysregulated DDR in a tumor cell creates genomic instability that is cancer enabling, and makes the tumor become dependent on alternative pathways such as base excision repair (BER) that may be targetable (e.g. PARP inhibitors). In this study we find evidence of dysregulated DDR by relative expression differences, as well as possible dysregulation to Mismatch Repair pathway. Several replication and repair genes were observed to have transcripts with aberrant splicing. This included CIZ1 which may be useful as an early biomarker, ATRIP a component of DDR, and MLH3 which may indicate abnormalities in mismatch repair. While there is some evidence that MMR and DDR are connected [10], we have examined and discuss the repair pathways separately. Interestingly, components of the replication machinery that lack mutation and SCNA correlate none-the less with increased expression, suggesting transcriptional mechanisms were used to overcome replication stress and fork collapse in UVM progression.

## Materials and Methods

### Mutation analysis

Mutational findings in this report are based upon data generated by the TCGA Research network and can be found at: http://cancergenome.nih.gov/ [11, 12].

### Relative Expression Analysis

RNA-seq–derived exon expression levels were first visualized in heat maps. The Gene Annotation File (GAF) “TCGA.hg19.June2011.gaf” (found originally at https://tcgadata.nci.nih.gov/docs/GAF/GAF.hg19.June2011.bundle/outputs/; now at https://gdc.cancer.gov/about-data/data-harmonization-and-generation/gdc-reference-files was used to create an exonStartStop.txt file for the gene(s) tested. This was used in turn to parse the “UVM.rnaseqv2__illuminahiseq_rnaseqv2__unc_edu__Level_3__exon_quantification__data.dat a.txt” file (originally at: https://confluence.broadinstitute.org/display/GDAC/Dashboard-Stddata; now at http://gdac.broadinstitute.org/) to create an “exonRPKM.txt” file used for standard Z score generation. Both files created, exonStartStop.txt and exonsRPKM.txt, were run through a verification step to confirm that the appropriate gene, TCGA barcodes, and RNAseq data were selected prior to their use. Exon start-stop sites from the exonStartStop.txt file were examined in Integrative Genome Viewer (IGV) [13, 14] using RNAseq data (https://portal.gdc.cancer.gov/) from the same case, to confirm the authenticity of the exon. The “Sashimi Plot” function in IGV was used to identify alternative splicing and isoforms from RNAseq data, with “Minimum junction coverage” routinely set at “4”.

In the “R” environment (https://cran.cnr.berkeley.edu/) Z scores were calculated for each exon of each gene by mean-centering, using the tumor cohort as average, the log2 transformed RPKM values and dividing by the standard deviation, visualizing high (red), no change/no expression (white), and low (blue) and arranging data by UVM cluster assignments (1–4) in heat maps.

### Placement of Cases into “High” and “Low” Expression Groups

An output file containing Z scores for each exon was created and used to calculate an average Z score for each gene. This was regarded reasonable as structural variations to genes were found only with low frequency. UVM cases were sorted into two groups, those with average Z scores “Above” and those “Below” zero. After group designation, the cluster (1–4) each case belonged to was identified and the numbers of cases “High” and “Low” for each cluster counted and displayed graphically using GraphPad Prism 6. Significant differences between “1 and 2” versus “3 and 4” were determined by Fisher’s exact (two tailed), using GraphPad Quick Calcs. Because very few genes and few cases (total n =80, in each cluster n=15 through 23) were examined, “q” values were not calculated, with the rationale that doing so might increase “type II” errors. Placement of cases into “high” and “low” grouping was made relative to the total tumor cohort average. Conjecturally, all cases “Below” the tumor cohort average could also be “Above” an average made from appropriate adjacent normal tissues, which was not available for RNA analysis in UVM samples (normal tissues for each UVM case were available for DNA analysis). The procedure was evaluated qualitatively using BAP 1 and RPS19 as test genes, and compared to mRNA differential expression Z scores generated by cBioPortal whose outcomes were made from RNAseq Version 2 using RSEM to perform quantitation, with the expression distribution of each gene compared only to tumors diploid for that gene when a “normal” cohort was not available.

### Clinical Data and Survival Analysis

The TCGA UVM cohort was made up of eighty matched tumor and normal specimens. Tumors were obtained from patients that did not have previous systemic chemotherapy or focal radiation, with appropriate consent obtained from institutional review boards. A panel of five histopathologists with expertise in ocular pathology and melanoma, examined hematoxylin and eosin stained sections from paraffin embedded tumors defining tumor extent, cell morphology, pigmentation, mitotic index, and the presence of tumor-infiltrating lymphocytes and macrophages. The following information was also curated: “Tumor status” (date of last contact), “Vital status” (dead/alive), “Date of last contact”, “Date of Death”, “Cause of Death”, “Other Cause of Death”, “New Tumor Event after Initial Treatment”, “Histology of New Tumor Event”, “Site of New Tumor”, “Other Site/new event”, “Date of New Event”, “Additional surgery”, “Additional Treatment/Radiation”, “Additional Treatment/Pharmaceutical”.

In principal four types of survival analysis can be made with TCGA clinical data. “Overall Survival” (OS) defined as the period from date of diagnosis until death from any cause, “Progression-Free Interval” (PFI) from date of diagnosis until the occurrence of an event in which the patient with or without the tumor does not get worse, “Disease-Free Interval” (DFI) date of diagnosis until first recurrence, and “Disease Specific Survival” (DSS) diagnosis date until death from the specific cancer type. All UVM Survival Curves constructed for this study were “Disease Specific Survival” curves, as recommended by TCGA Pan Cancer Guidelines https://wiki.nci.nih.gov/display/TCGAM/2017-03-23+PCA+Network. Kaplan Meier survival plots were constructed using GraphPad Prism 6.0 software. The Log-rank (Mantel-Cox) and Hazard Ratio tests were used to determine significance.

## Results

### BAP1 and RPS19

A total of thirty-seven genes were studied (Table 1). Mutations and differential expression for the gene set have been depicted in an “Oncoprint” from cBioPortal (http://www.cbioportal.org/public-portal) (Figure 1). In this study we examine expression by two algorithms, and use the terms “differential” versus “relative” to distinguish between them. “Differential” expression is defined as being made with respect to a non-neoplastic or, as is the case for UVM, an estimated normal sample (see Materials and Methods). “Relative” expression is specified as comparison to the total UVM tumor cohort average and is made for both mRNA and individual exons.

**Table 1.**
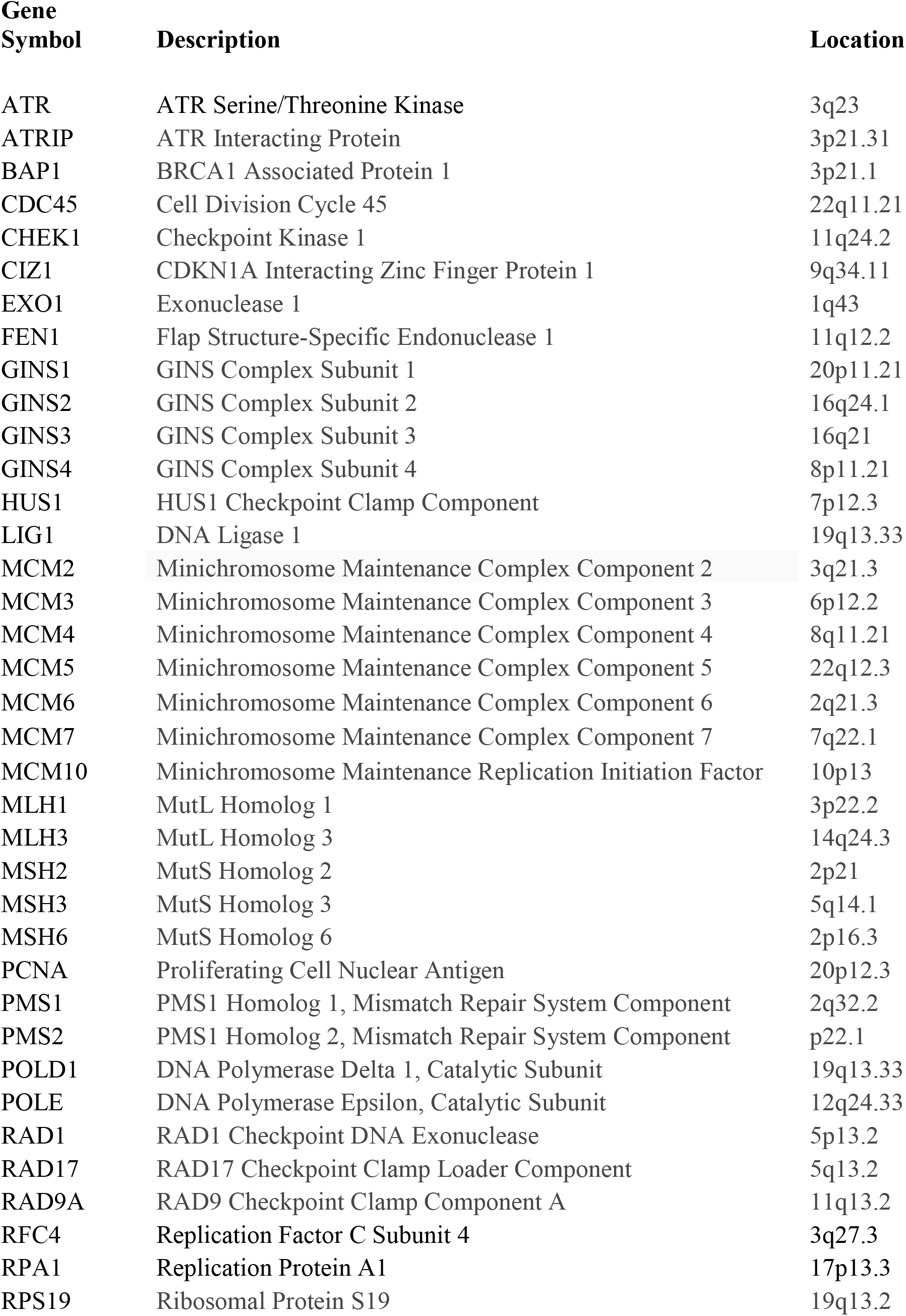
A Selected List of Genes Associated with Replication and DNA Repair

**Figure 1.**
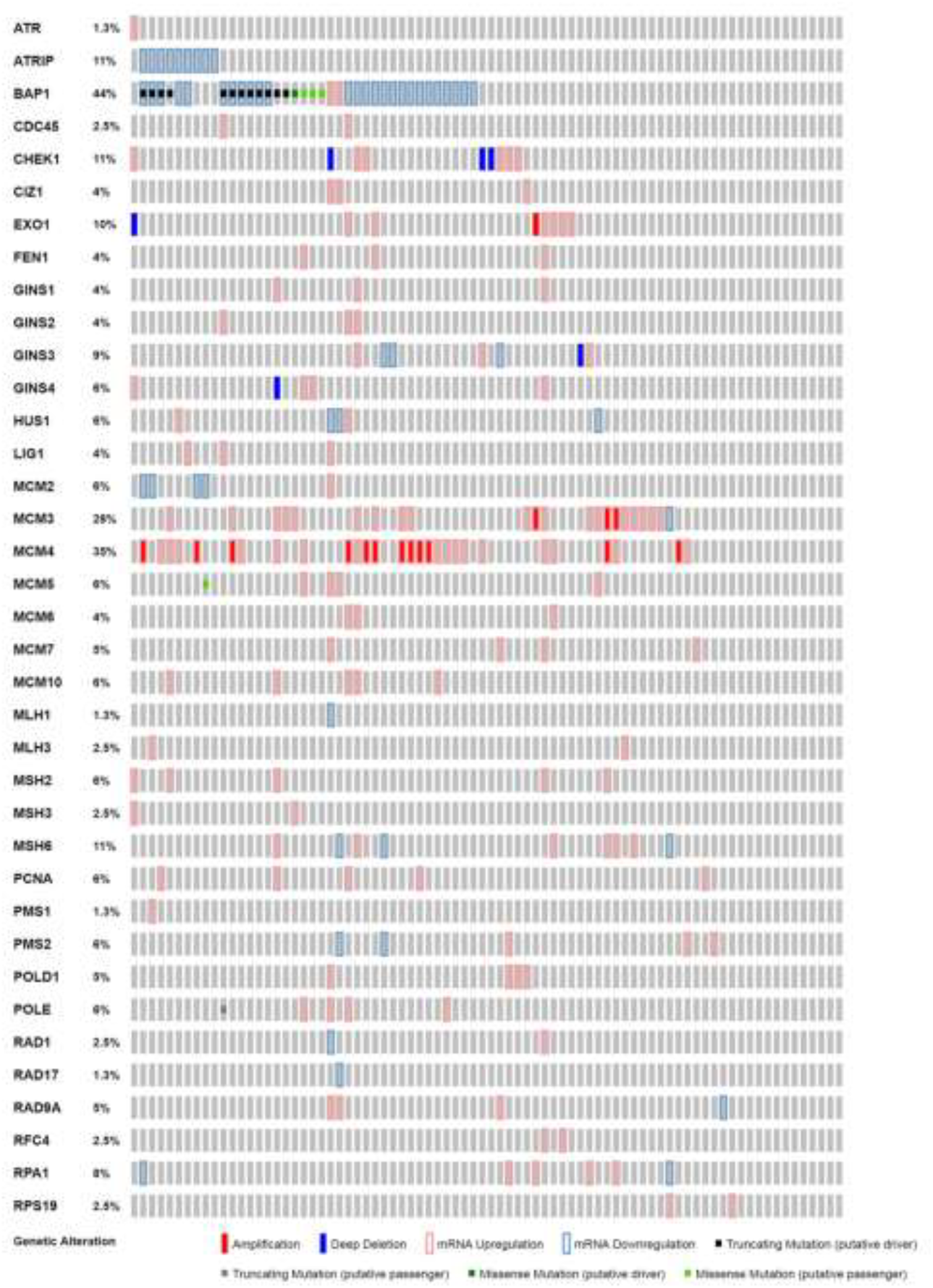
Oncoprint Displaying Mutations and differential expression of Selected Genes in Replication and Repair.

BRCA 1 Associated Protein 1 (BAP 1) and RPS19 were used to compare the methods of analysis (Table 2). Examination of BAP 1 differential expression Z scores calculated by cBioPortal, showed a two-fold greater inclusion of cases “Below” average expression. For this particular tumor type due to the relationship of monosomy 3 to subtypes and survival, the approach using an estimated reference alters comparison to relative expression for genes located on chromosome 3. SCNA subtypes 3 and 4 are both monosomic for chromosome 3 and constitute approximately half the total tumor cohort. In contrast using RPS19 as a test gene, a gene which codes for a 40S Ribosome complex protein with cytological location at chromosome 19q13.2, showed no significant difference in relative expression found between the study method and cBioPortal values. UVM do not have significant SCNA for chromosome 19, the four additional cases found in the “Below” group of cluster 3 are due to the use of RSEM verses RPKM. These results show incongruity between differential expression about an estimated normal value and relative expression about a tumor cohort average, when high numbers of cases are not diploid. We would like to note explicitly that presentation of the discrepancy is not meant to claim one set of calculations superior to the other, but to explain why additional calculations were made for relative expression.

**Table 2.**
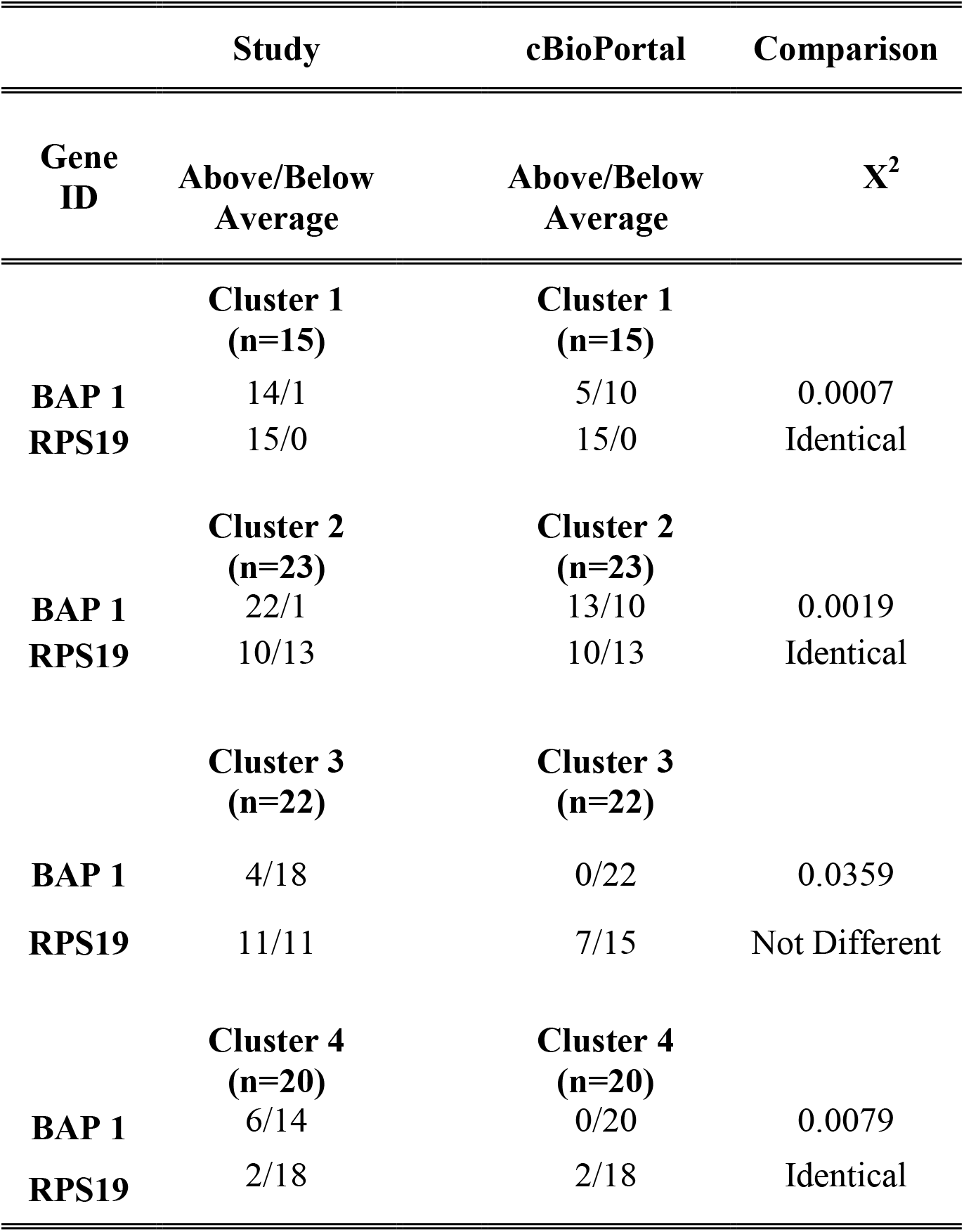
Comparison Study vs cBioPortal

|  | **Study** | **cBioPortal** | **Comparison** |
| --- | --- | --- | --- |
| **Gene ID** | **Above/Below Average** | **Above/Below Average** | **X<sup>2</sup>** |
|  | **Cluster 1**<br>**(n=15)** | **Cluster 1**<br>**(n=15)** |  |
| **BAP 1** | 14/1 | 5/10 | 0.0007 |
| **RPS19** | 15/0 | 15/0 | Identical |
|  | **Cluster 2**<br>**(n=23)** | **Cluster 2**<br>**(n=23)** |  |
| **BAP 1** | 22/1 | 13/10 | 0.0019 |
| **RPS19** | 10/13 | 10/13 | Identical |
|  | **Cluster 3**<br>**(n=22)** | **Cluster 3**<br>**(n=22)** |  |
| **BAP 1** | 4/18 | 0/22 | 0.0359 |
| **RPS19** | 11/11 | 7/15 | Not Different |
|  | **Cluster 4**<br>**(n=20)** | **Cluster 4**<br>**(n=20)** |  |
| **BAP 1** | 6/14 | 0/20 | 0.0079 |
| **RPS19** | 2/18 | 2/18 | Identical |

### Pre-Replication and Pre-Initiation Complexes

The Pre-Replication, Pre-Initiation, and Replisome complexes, DDR and MMR pathways have been represented schematically to indicate where the selected components of this study function (Figures 2–4). Examination of differential expression plots across tumor types for the Mini-Chromosome Maintenance (MCM) helicase components of the Pre-Replication Complex show increased expression above “normal” for all components. MCM2 (3q21.3) exemplifies this general trend (Figure 5) (http://gdac.broadinstitute.org/runs/stddata__2016_01_28; doi:10.7908/C11G0KM9). Relative expression profiles show MCM2 drops below the average significantly in UVM clusters 3 and 4 (P= 0.0001) (Table 3), correlating with increased malignancy and decreased disease specific survival (P= 0.0001). Four of these cases can be seen using differential expression (Figure 1). Half of the MCM2-7 complex components are located on chromosomes that the TCGA finds by SNP-based Copy Number Analysis to have copy number alterations that include monosomy chromosome 3, 8q and 6p gains. Comparing relative expression for all genes in this study found on chromosome 3 indicates relative expression levels do not always simply correlate with SCNA (Figure 6A-B), reflecting the TCGA finding that expression subtypes are only partially concordant with SCNA subtypes [1].

**Table 3.**
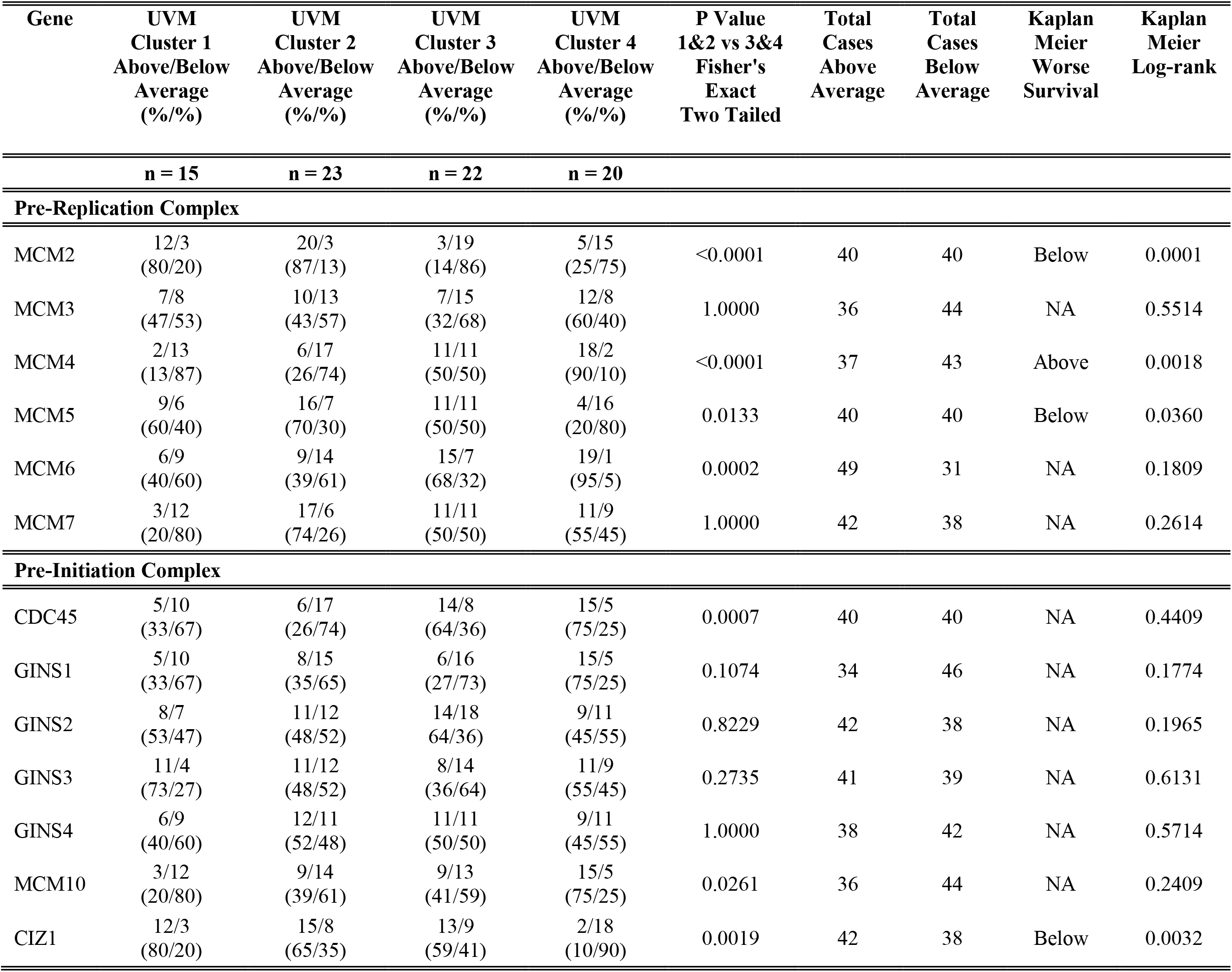
Relative Expression and Survival Correlation of Pre-Replication and Pre-Initiation Complex Factors

**Figure 2.**
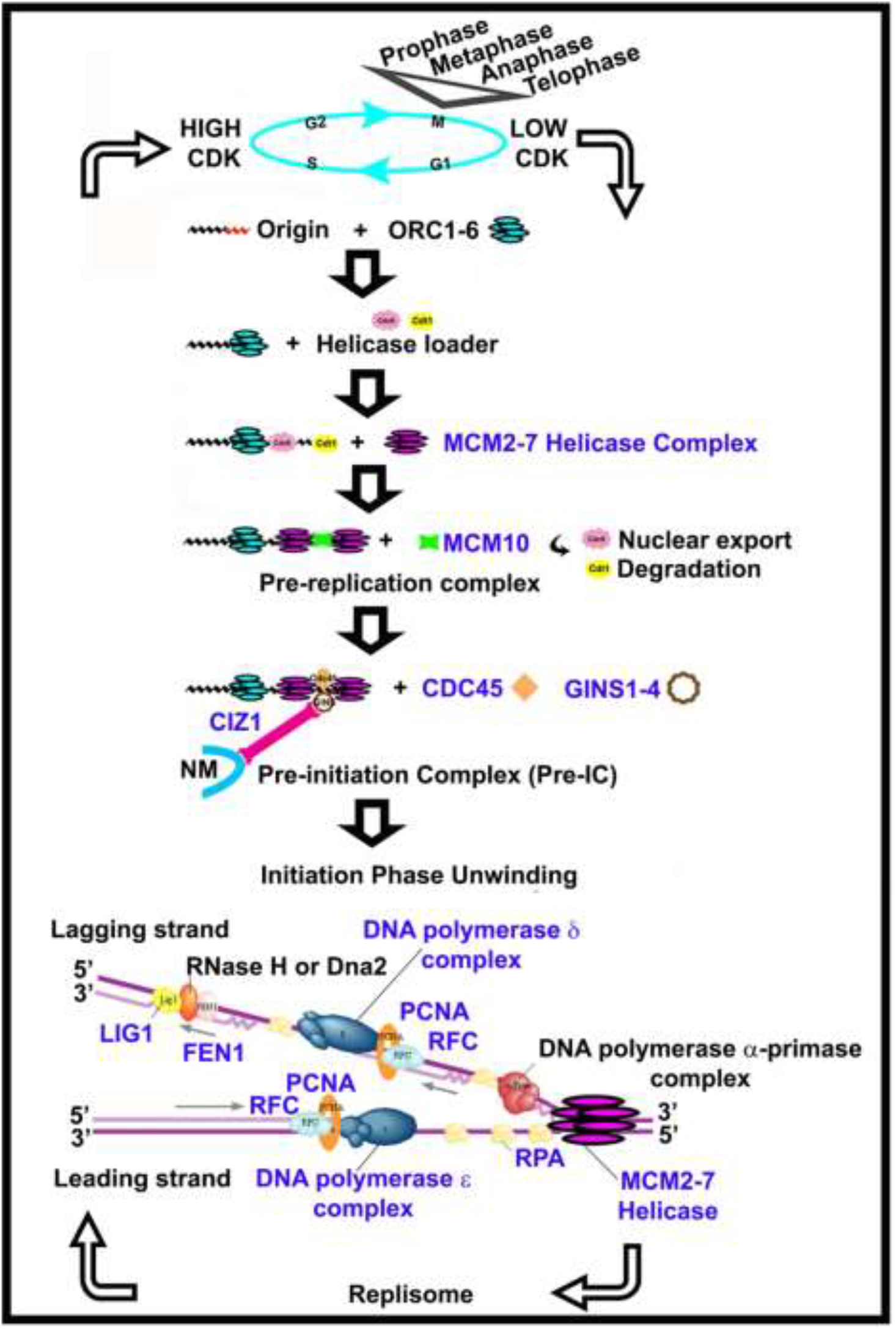
Schematic Representation Depicting Selected genes in the context of Replication. Selected genes in replication (*blue*) (Replisome from http://www.genome.jp/kegg/pathway.html).

**Figure 3.**
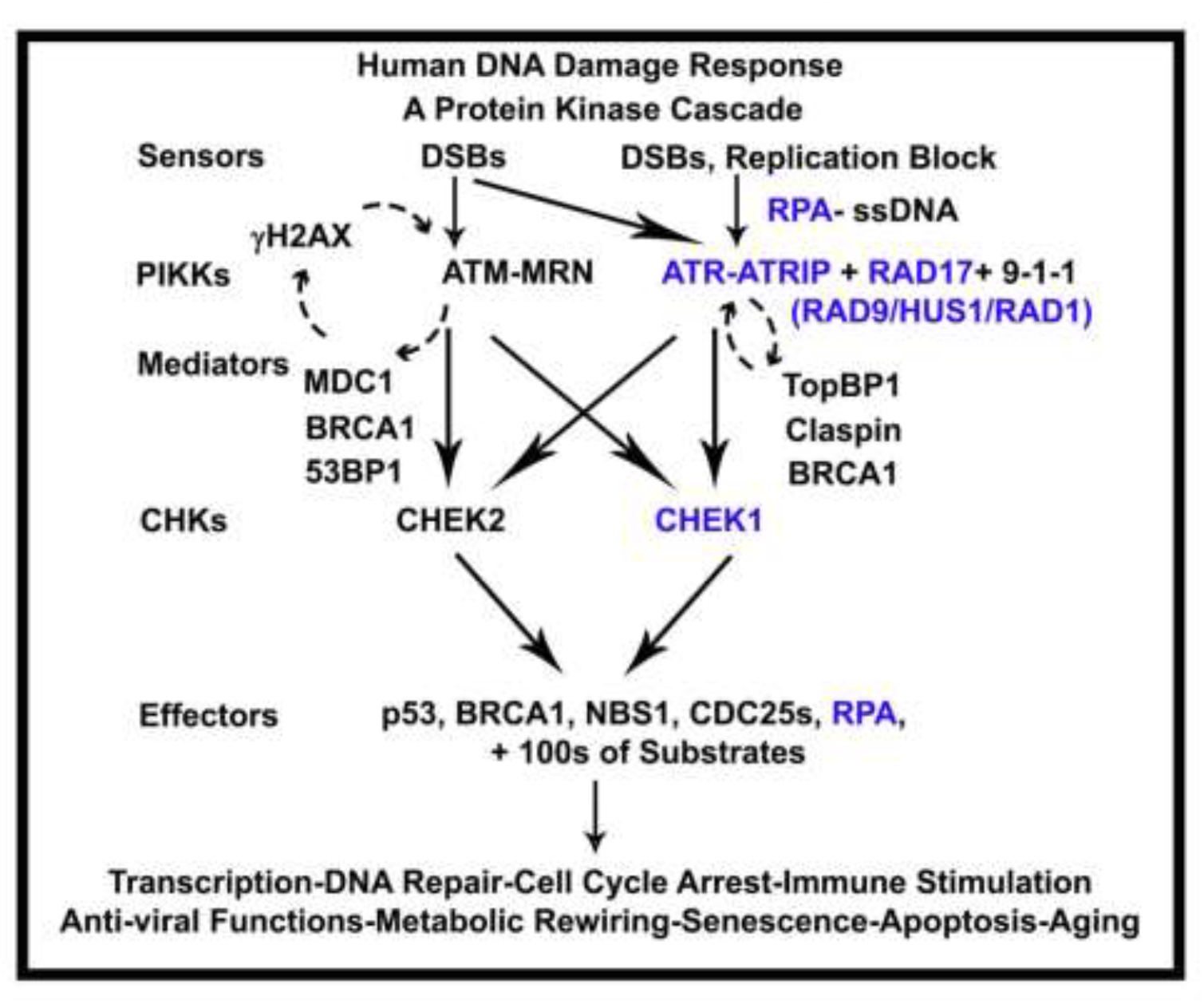
DNA Damage Response (DDR). Selected genes in replication (*blue*), (diagram from http://elledgelab.med.harvard.edu/?page_id=264).

**Figure 4.**
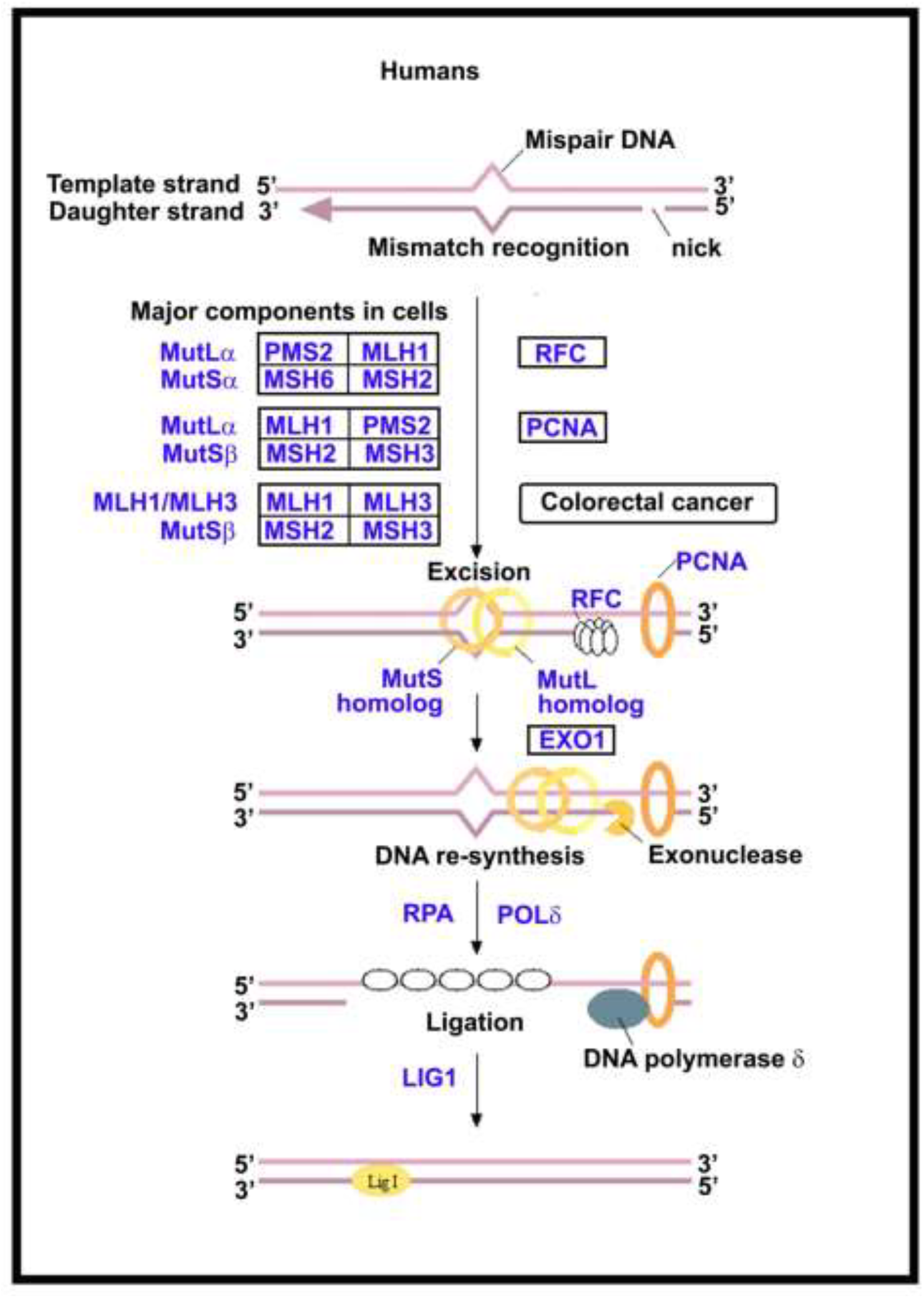
Mismatch Repair (MMR). Selected genes in replication (*blue*), (diagram from http://www.genome.jp/kegg/pathway.html).

**Figure 5.**
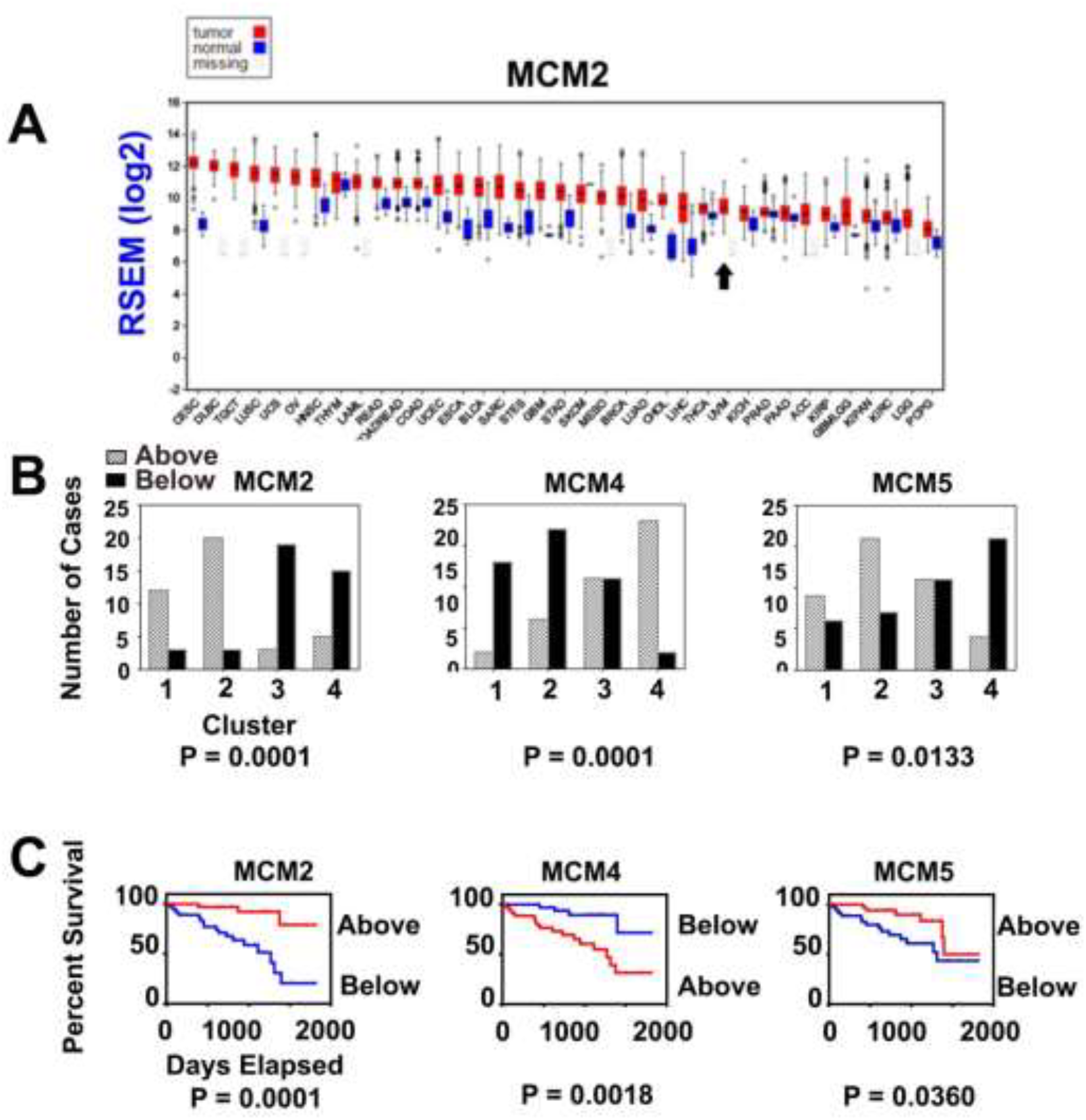
Changes to the Mini Chromosome Maintenance (MCM) Helicase Complex (A) Differential expression of MCM2 across tumor types, highest to lowest expression UVM (black arrow). (B) Number of cases in cluster [1–4] with Z scores “above” (hatched) and “below” (black) tumor cohort average, Fisher’s Exact test, two tailed. (C) Kaplan Meier Survival Plot, “above” average (red) “below” average (blue), Log-rank test.

**Figure 6.**
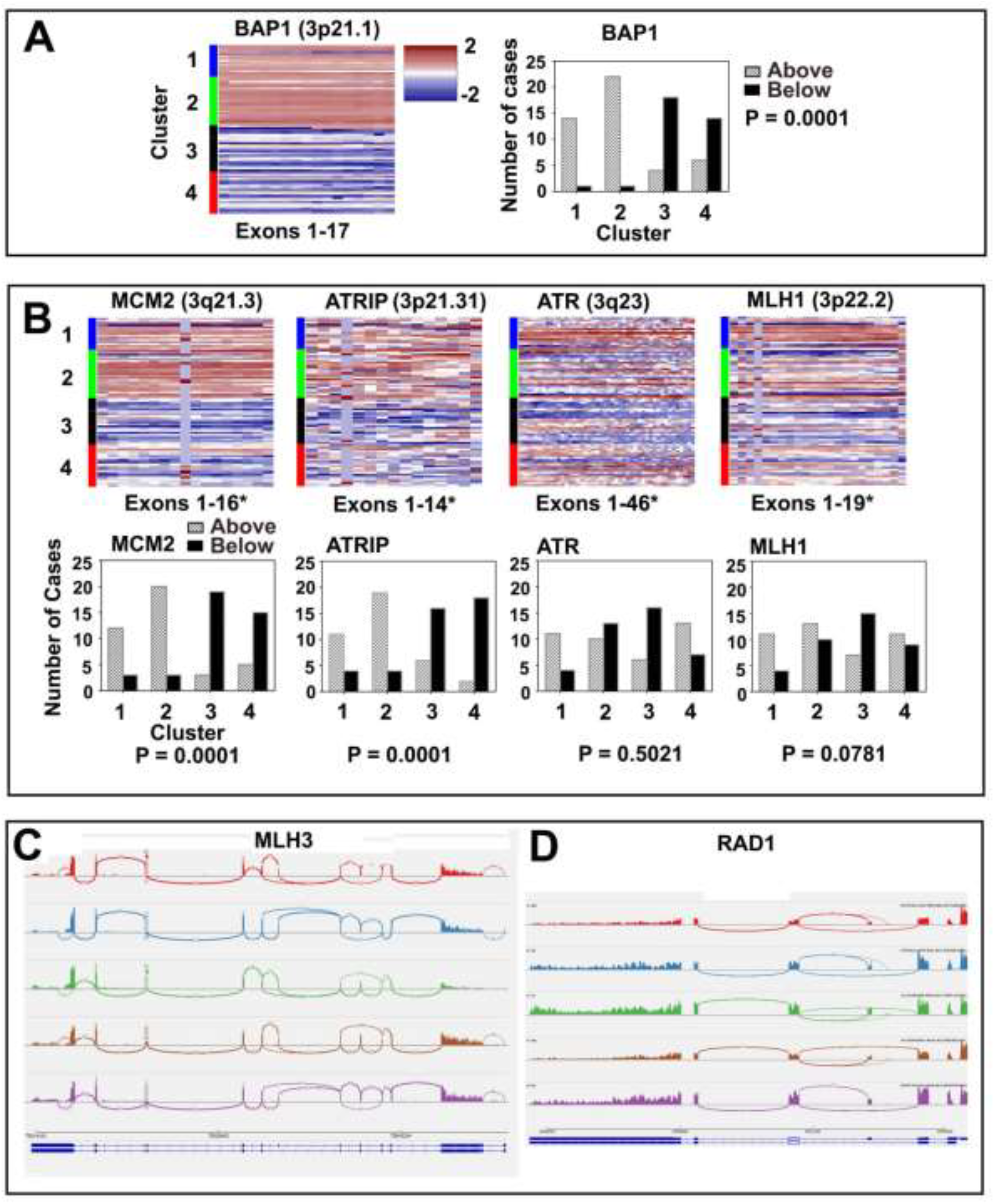
Comparison of Chromosome 3 Genes. (A) BAP1 relative expression (*left*), number of cases in cluster (*right*), 1 and 2 versus 3 and 4 (Fisher’s Exact test, two-tailed). (B) MCM2, ATRIP, ATR, MLH1. (C) Sashimi plot (IGV), MLH3. (D) Sashimi plot, RAD1.

MCM4 (8q11.21) has increased relative expression in clusters 3 and 4, with increased expression having worse survival that correlate with CNV gains to chromosome 8q in cluster 4 (P= 0.0018) (Figure 5, Table 3). MCM5 (22q12.3) had lower relative expression in UVM clusters 3 and 4, with cases having worse survival. For MCM5, the P values are significant but less convincing (P = 0.036), (see supplemental Figure 1 for data covering all MCM helicase components).

We include CDC45, GINS1-4, MCM10, and CIZ1 in the Pre-replication complex (see discussion). While CDC45 and MCM10 appeared highly expressed in the higher risk subtypes, alterations did not correlate with survivability (Table 3). GINS 1–4 were not differentially expressed across the clusters. CIZ1 (Cip1-interacting zinc finger protein 1) [15–20], had highly significant difference between the clusters, with lower expression correlating to higher risk subtypes and decreased survival (P = 0.0019) (Table 3, Figure 7 B-D). We also found evidence of exon skipping in exon 4 throughout the tumor cohort previously seen in Ewing tumor [21] and Lung Cancer [22] and the C terminal region (Figure 7 B & E), both by relative expression and by examining mRNA isoforms using the Sashimi Plot function in IGV (Figure 7 E).

**Figure 7.**
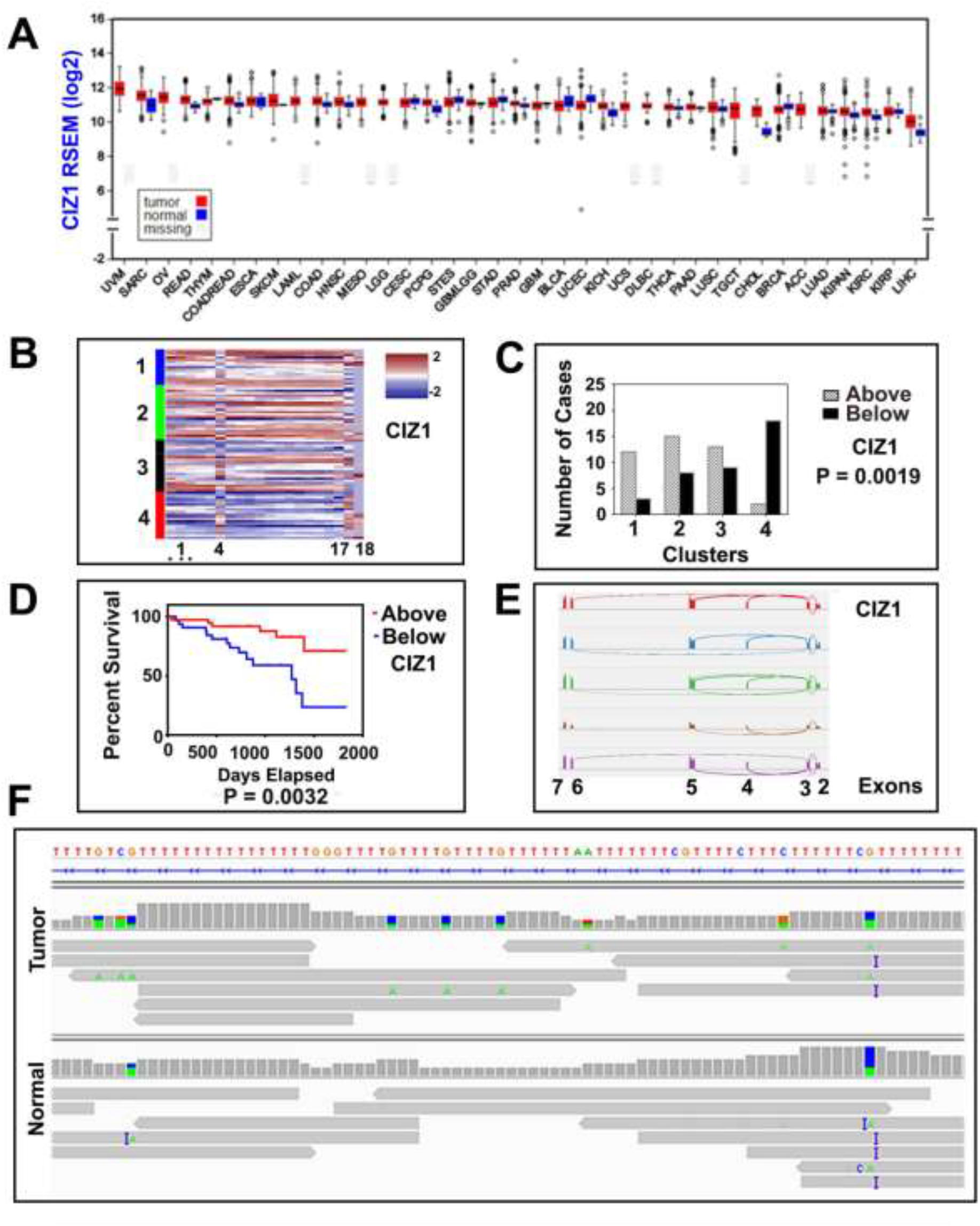
CIZ1 (A) Differential Expression, across tumor types. (B) CIZ1 relative expression, exon1 alternative splicing (asterisk). (C) Number of cases, 1 and 2 versus 3 and 4 (Fisher’s Exact test, two-tailed). (D) Kaplan Meier Survival Plot (Log-rank test). (E) Sashimi Plot (IGV), exon 4 skipping. (F) CIZ1 Intron 3 mononucleotide repeat (hg19:130,950,210-130,950,372), adenine insertions (purple bar).

A mononucleotide repeat in CIZ1 intron 3 was previously suggested to explain exon 4 skipping mechanistically, and hypothesized to be the result of MMR deficiency [21]. Fifty out of 80 UVM tumors did have low pass WGS available for tumor and normal counterpart tissues as well as tumor RNAseq, and we examined them for alteration in CIZ1 Intron 3. Viewing the mononucleotide repeat (hg19:130,950,210-130,950,372) in IGV from these fifty tumor/normal pairs showed 94% of the tumors and 92% of the normal samples had some mononucleotide repeat alteration (single nucleotide alteration included) (Figure 7 F). We could not clearly discern MSI from possible sequencing artefacts that are suggested by the high alteration frequency. These results and MMR status in relation to CIZ1 mononucleotide repeat alteration are discussed more fully below.

### Replisome, DNA Damage Response, Mismatch Repair Proteins

In addition to alterations in the members of the MCM complex in UVM, there is evidence of transcriptional upregulation to integral components of the replisome, and to components of DNA repair pathways. Sliding clamp component Proliferating nuclear cell antigen (PCNA) and Flap endonuclease (FEN1), well known markers of leading and lagging strand synthesis, were found to have increased expression across clusters, as did LIG1 and the DDR clamp component HUS1 (Figure 8 A and B). PCNA and HUS1 increase correlated significantly with worse survival, FEN1 and LIG1 did not (Table 4). PCNA had noticeable expression differences to Exon 6 in comparison to other exons however splicing alterations were not identified in IGV. Other replication components RFC4, responsible for elongation of primed DNA templates, and RPA1, a stabilizer of single stranded DNA, did not show differences across the clusters (data not shown).

**Table 4.** Relative Expression and Survival Correlation of Replisome

| Gene | UVM<br>Cluster 1<br>Above/Below<br>Average<br>(%/%) | UVM<br>Cluster 2<br>Above/Below<br>Average<br>(%/%) | UVM<br>Cluster 3<br>Above/Below<br>Average<br>(%/%) | UVM<br>Cluster 4<br>Above/Below<br>Average<br>(%/%) | P Value<br>1&2 vs<br>3&4<br>Fisher's<br>Exact<br>Two<br>Tailed | Total<br>Cases<br>Above<br>Average | Total<br>Cases<br>Below<br>Average | Kaplan<br>Meier<br>Worse<br>Survival | Kaplan<br>Meier<br>Log-rank |
| --- | --- | --- | --- | --- | --- | --- | --- | --- | --- |
|  | n = 15 | n = 23 | n = 22 | n = 20 |  |  |  |  |  |
| **Replisome** |  |  |  |  |  |  |  |  |  |
| PCNA | 8/7<br>(40/60) | 4/19<br>(17/83) | 8/14<br>(36/64) | 17/3<br>(85/15) | 0.0147 | 37 | 43 | Above | 0.0280 |
| FEN1 | 5/10<br>(33/67) | 9/14<br>(39/61) | 11/11<br>(50/50) | 17/3<br>(85/15) | 0.0132 | 42 | 38 | Above | 0.0884 |
| LIG1 | 1/14<br>(7/93) | 10/13<br>(44/56) | 16/6<br>(73/27) | 15/5<br>(75/25) | 0.0001 | 42 | 38 | NA | 0.2590 |
| POLD1 | 7/8<br>(47/53) | 17/6<br>(74/26) | 7/15<br>(32/68) | 6/14<br>(30/70) | 0.0067 | 27 | 43 | Below | 0.0597 |
| POLE | 4/11<br>(27/73) | 14/9<br>(61/39) | 11/11<br>(50/50) | 16/4<br>(80/20) | 0.1761 | 45 | 35 | Above | 0.0990 |
| **DNA Damage Response** |  |  |  |  |  |  |  |  |  |
| HUS1 | 7/8<br>(47/53) | 9/14<br>(39/61) | 12/10<br>(55/45) | 18/2<br>(90/10) | 0.0124 | 46 | 34 | Above | 0.0278 |
| RAD9A | 9/6<br>(60/40) | 10/13<br>(43/57) | 13/9<br>(59/41) | 16/4<br>(80/20) | 0.1105 | 48 | 32 | NA | 0.8086 |
| RAD1 | 8/7<br>(53/47) | 10/13<br>(43/57) | 10/12<br>(45/55) | 16/4<br>(80/20) | 0.2609 | 44 | 36 | NA | 0.1076 |
| RAD17 | 8/7<br>(53/47) | 9/14<br>(39/61) | 9/13<br>(41/59) | 16/4<br>(80/20) | 0.2624 | 42 | 38 | NA | 0.5653 |
| ATR | 11/4<br>(73/27) | 10/13<br>(44/56) | 6/16<br>(27/73) | 13/7<br>(65/35) | 0.5021 | 40 | 40 | NA | 0.8707 |
| CHEK1 | 4/11<br>(27/73) | 8/15<br>(35/65) | 12/10<br>(55/45) | 15/5<br>(75/25) | 0.0041 | 39 | 41 | NA | 0.3072 |
| ATRIP | 11/4<br>(73/27) | 19/4<br>(83/17) | 6/16<br>(27/73) | 2/18<br>(10/90) | 0.0001 | 38 | 42 | Below | 0.0001 |

**Figure 8.**
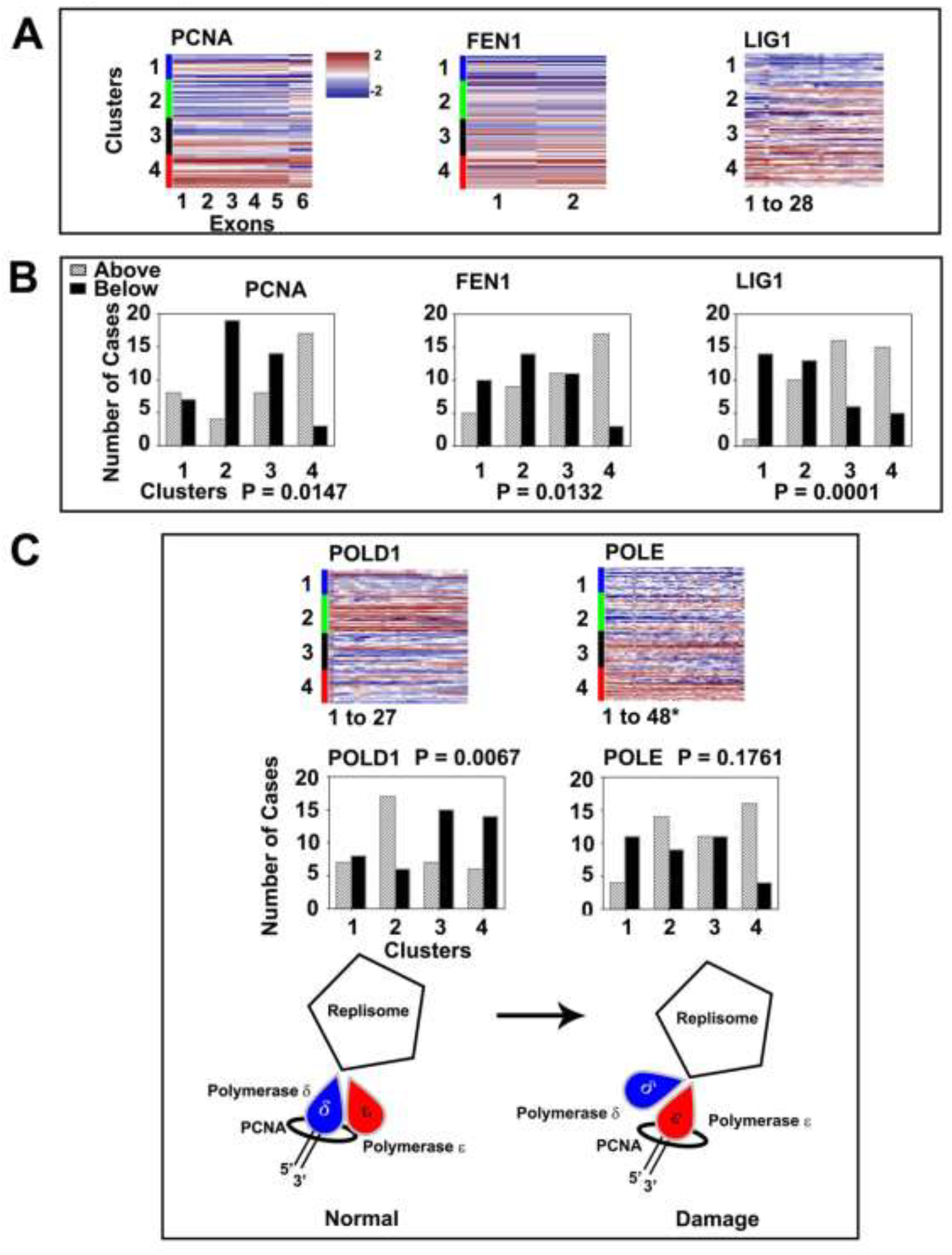
Replisome Components (A) Proliferating Nuclear Cell Antigen (PCNA), Flap endonuclease (FEN1), and Ligase 1 (LIG1), Relative Expression. (B) Number of cases PCNA, FEN1, LIG1, POLD1, POLE (Fisher’s exact test, two tailed. (B) Simplified model of switch to POLE upon DNA damage.

Two polymerases were examined for relative expression, POLD1 and POLE. POLD1 relative expression was significantly different in clusters 1 and 2 compared to 3 and 4, survival correlation was not (P = 0.06, Log rank). POLE expression was not significantly different across clusters 1 through 4. Examining POLD1 and POLE clusters 3 & 4 to each other however, showed significant difference in the number of cases “above” and “below” the mean (P = 0.0042 by Fisher’s exact test, two tailed), indicating clusters 3 and 4 had more cases with higher than average POLE expression than POLD1 (Figure 8 C).

Upon DNA damage the 9-1-1 response is activated to stop cell cycle progression (Figure 3). HUS1 is a member of this pathway that acts as part of the replication “sliding clamp”. Its relative expression was significantly increased across the UVM clusters (Table 4). RAD1, RAD9A, and RAD17, also members of the DDR complex, did not have significant expression or survival differences. RAD1 showed evidence of alternative splicing involving exon 3 previously reported as a natural isoform (Figure 6D).

MLH3 and MSH6 transcripts had significant expression differences between clusters 1 and 2 versus 3 and 4 however there were no survival advantages or disadvantages (Table 5). The MLH1 relative expression pattern across UVM SCNA subtypes did not completely behave as anticipated from simple correlation with CNV (Figure 6). Because MSI was reported in CIZ1 intron 3, we examined UVM RNA-seq data for several MMR genes in IGV and found further evidence of exon skipping and alternative splicing. MLH3, a protein that directly interacts with MLH1, is thought to repair DNA insertion/deletion loops. Mutations to MLH3 result in MSI of short repetitive sequence length instability, similar to that found in CIZ1 intron 3 [23]. Examination of UVM RNA-seq data for MLH3 led to the identification of an isoform by Sashimi plots with deletions in both exons 7 and exon 5 (Figure 6C) simultaneously. MSH6 was also examined in IGV for alternative splicing, which was not observed.

**Table 5.**
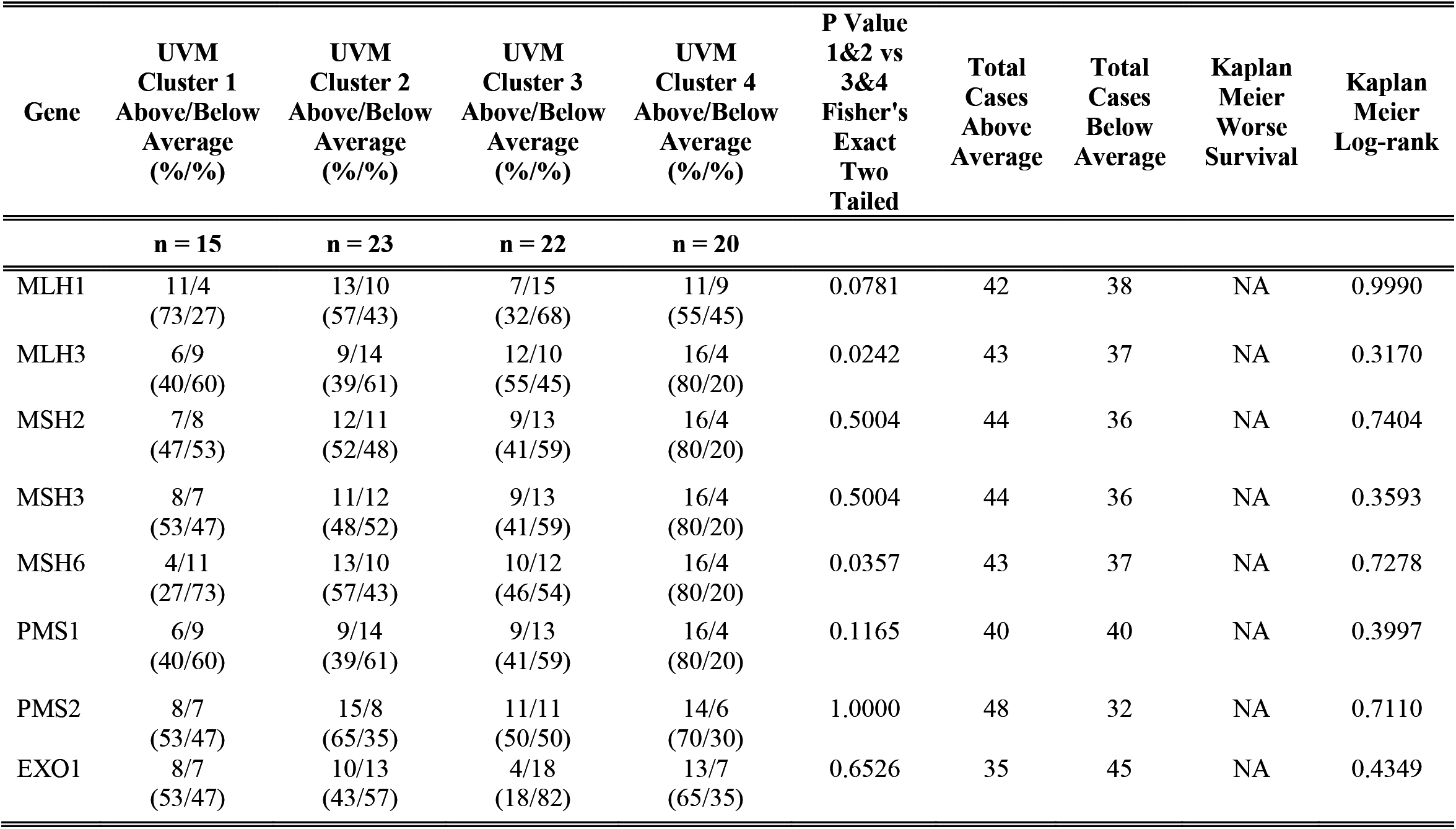
Relative Expression and Survival Correlation of Mismatch Repair Genes

## Discussion

In this report, selected components of replication and repair processes of UVM were examined for transcriptional alteration with or without gene mutation. Two approaches were taken for evaluating transcriptional changes that permitted distinct vantage points for data interpretation. In the first method, tumor mRNA levels were compared to normal (estimated in UVM) mRNA. In the second method, tumor mRNA levels were compared to the tumor cohort average (actual) at both mRNA and exon levels permitting comparison across subtypes identified by TCGA on the basis of SCNA.

Relative expression differences were found across the UVM cohort in components of the pre-replication and pre-initiation complexes. These include the Mini Chromosome Maintenance helicase complex (MCM2-7) components that effectually act to “license” the firing of origins of replication during G1-phase of the cell cycle, determining the multiple nascent origins that replicate during S phase. A recent comprehensive review of the MCM2-7 helicase proteins has been made by Riera et al. [24]. The complex is thought to load in excess, with dormant origins becoming active when nearby replication forks have stalled functionally serving as backup. Mouse models deficient in MCM components have high rates of cancer and genome instability [25–29], which supports the concept of relative deficiency of MCM factors in UVM. Decrease in MCM2 specifically is thought to reduce replication initiation in gene rich early replicating regions, and to increase genetic damage at a subset of gene rich locations [30] without changing the replication rate. The eukaryotic replisome *in vitro* has only been reconstituted at a rate four times lower than that *in vivo*, implying that not all factors contributing to replication rate are known [31]. That individual component increase or decrease may have implications for replication rate while possible, is not verifiable from current data. Another putative function for MCM2 due to its histone binding and chaperone capacity may be to orchestrate histone dynamics during replication [32], and it has also been linked conceptually to transcription [33]. UVM appear to have MCM concentration alterations including decreased MCM2, increased MCM4 and decreased MCM5 that correlate with increased malignancy and decreased survival. Overexpression of MCM4 has been found previously in multiple transformed cell lines and numerous tumor types [34–36]. The unbalanced expression of MCM2-7 individual components likely affects the quantitative amount of functional helicase available, lowering the ability to load in excess and contributing to unregulated replication.

We discuss CIZ1 in the context of the pre-initiation complex. Its inclusion is relatively recent [20] and less well documented. The gene plays a role in cell cycle control and replication by interaction with Cip1/Waf1[15], and coordinating cyclin E and A functions in the nucleus [17]. It is required for cells to enter into S phase [20]. CIZ1 physically tethers to non-chromatin nuclear structures in the nuclear matrix, and co-localizes in immune-histochemical studies with PCNA [15]. Two functionally distinct domains exist, an N-terminal domain involved in replication and a C-terminal domain comprising the nuclear matrix anchor [16]. Depletion of transcripts decreases proliferation *in vitro* [15]. Alternatively spliced variants are found in cancers and other disorders that appear to alter protein localization [19]. Variant CIZ1 has recently been published as a circulating biomarker for early-stage lung cancer [22]. Differential expression plots for CIZ1 show the gene is most highly expressed in UVM. Relative expression across the TCGA SCNA subtypes shows clusters 1, 2, and 3 have CIZ1 expression above the cohort average and cases below the tumor average are primarily in subtype 4. Mutations to splicing factors SF3B1 and SRSF2 have been found in UVM subtypes 2 and 3. Spliceosome alterations are hypothesized to “drive” the tumor type. Almost all UVM cases examined show evidence of exon 4 alteration previously reported in other cancers as well as alteration to the carboxyl end of the protein thought to be responsible for nuclear localization. Further study of CIZ1 transcripts in relation to spliceosome alterations and examination of the sera of UVM patients for variant CIZ1 transcripts may yield an early UVM biomarker.

The findings for CIZ1 led to identification a mononucleotide repeat in CIZ1 Intron 3 with possible MSI, and as a result we re-examined MMR for alteration. Almost no evidence points to the involvement of MMR components as drivers for UVM. With the exception of EXO1, there are no mutations to MMR genes, no significant differential expression, and no survival disadvantage from relative expression alteration. Examination of mutation rate does not show logarithmic elevation, typical of MMR deficient tumors. Despite location on chromosome 3, MLH1 relative expression did not correlate with SCNA. However, differences in relative expression across the UVM clusters were found for MLH3 and MSH6. We also found evidence (*prima facie*) of exon skipping in MLH3 (MLH3Δexon5/Δexon7), a protein that interacts with MLH1 directly. MLH3Δexon7 has been previously described [23] and in that study the isoform lacked MLH1 binding activity. Fifty-five percent of UVM cases with WGS had evidence of the MLH3Δexon5/Δexon7 isoform, suggesting a mechanism for partial MMR defect in UVM and during development with resulting predisposition. In addition to playing a role in MMR, MLH3 also has a meiotic phenotype. Evidence of MSI was observed (CIZ1 mononucleotide repeat in Intron 3) in both tumor tissue and its normal counterpart. Mononucleotide tracks are difficult to sequence and the frequency of MSI found is implausibly high, however, some examples genuinely appear have alterations in the germline that also appear in the tumor (Figure 7F). Interestingly, examination of differential expression across tumor types (Figure 7A) shows CIZ1 not much different between tumors and their normal tissue quantitatively, unlike MCM2 for example (Figure 5A).

Reviewing relative expression for PCNA, FEN1, and LIG1 we found increase that implied their involvement in overcoming the replication stress evidenced by factors in the pre-replication and pre-initiation complexes. PCNA is a ring shaped homo-trimer that encircles DNA [37]. It interacts competitively with many other factors at the PIP motif [38] [39] and is an essential co-factor for DNA polymerases during replication. It is involved in repair processes and can also be modified post-translationally by phosphorylation, ubiquitination, SUMO proteins, ISGylation, Acetylation, and S-nitrosylation. Each modification has corresponding biological functions that include proliferation, MMR inhibition, translesion synthesis, homologous recombination, genomic stability, and apoptosis, respectively. One of its major roles is to promote tolerance to DNA damage during replication [40]. Because of its role in proliferation, PCNA is a target for cancer therapy. Several small molecules that either block PCNA-Polδ or PCNA trimer formation have been identified with proliferation-inhibitory effects in vitro [37] [41–43].

Polymerase processivity is activated by PCNA. Two polymerases, POLD1 and POLE, important in the replication of B form DNA were selected for observation, as examination of polymerase behavior it its entirety was beyond the scope of this study. POLD1 relative expression was decreased in subtypes 3 and 4, POLE was increased. The expression levels of POLD1 and POLE support a recently proposed model [44] in which a switch to POL ε and away from POL δ occurs upon DNA damage. We have not yet addressed the behavior of POL α, or of the specialized polymerases that recently have been hypothesized to assist the replication machinery in the prevention of replication stress [4].

In our examination of DDR components we saw significant decrease in relative expression of ATRIP (ATR Interacting Protein) in clusters 3 and 4 [45–49]. Nine of these cases (11% of UVM) were also found by differential expression (Figure 1). Exon 3 deletion was identified in ATRIP transcripts across the UVM subtypes (Figure 4). PARADIGM and MARINa algorithms previously used to examine the DDR pathway did not find DDR impairment. ATRIP is located at 3p21.31 (very close to BAP1, 3p21.1), suggesting the disparity might be a result of using an estimated normal reference value. Interestingly as mentioned in the TCGA study [1], recent reports suggest that BAP1 itself may function in DDR [50] [51].

## Conclusions

In summary, we have examined selected replication and repair factors in UVM with known biology for expression differences with and without mutation across TCGA SCNA subtypes. We have observed differences in expression that support the concept that these genes and their products are causally involved in the origin and progression of uveal melanoma. These data also suggest further avenues of research for biomarker identification and therapeutic approach. The few genes examined in this study are prototypical for the kind of findings that can be made examining RNAseq data using relative expression. We are currently modifying the methodology to be more inclusive, routine, and high throughput to rapidly screen all replication and repair genes across all TCGA tumor types.

## Abbreviations

TCGA: The Cancer Genome Atlas
UVM: Uveal Melanoma
WES: Whole exome sequencing
WGS: Whole genome sequencing
SCNA: Somatic copy number alterations
DDR: DNA Damage Repair
MMR: Mismatch Repair
BER: Base excision repair
MCM: Mini-Chromosome Maintenance
RPKM: Reads per Kilobase of transcript per million mapped reads
RSEM: RNAseq by Expectation Maximization

## Declarations

Ethics approval and consent to participate were initially obtained by TCGA; all data are now in the public domain. Links to the data repositories have been given in the manuscript. The author consents for publication by BMC Cancer, and declares no conflict of interest. The study was supported by a grant from the National Institutes of Health to Raju Kucherlapati for the Harvard Genome Characterization Center No. 5U24CA144025.

## Acknowledgements

We thank Dr. Raju Kucherlapati and Dr. A. Gordon Robertson for reading the manuscript and providing comments and advice. Relative expression was determined by modifications to a method created for the analysis of structural fusions by Dr. Christopher Bristow. The work was also supported by Research Computing at Harvard Medical School, and we specifically thank Ms. Kristine Holton and Mr. Alex Truong for their very helpful consultation.

---

**Supplemental Figure 1** Changes in relative expression to total components of the Mini Chromosome Maintenance (MCM) Helicase Complex.

